# Loss of *CDKN2A* remodels Golgi morphology to create a lipid peroxidation vulnerability

**DOI:** 10.64898/2026.11.10.765755

**Authors:** Amandine Amalric, Baixue Yang, Rebecca Scarpati, Apoorva Uboveja, Ruiying Zheng, Alex Tom, Aidan R. Cole, Andrea Andress Huacachino, Qin Liu, Nathaniel W. Snyder, Andrew V. Kossenkov, Katherine M. Aird, Raquel Buj

## Abstract

Loss of the tumor suppressor *CDKN2A*/p16 occurs in ∼50% of human cancers, yet no therapies specifically target tumors with complete loss or reduced *CDKN2A*/p16 expression (hereafter *CDKN2A*/p16^Low^). Identifying selective vulnerabilities in *CDKN2A*/p16^Low^ tumors therefore represents a critical unmet need. Here, we identify altered Golgi morphology as a vulnerability in *CDKN2A*/p16^Low^ melanoma. A CRISPR knockout screen revealed that genes associated with Golgi membranes are essential for *CDKN2A*/p16^Low^ cell survival. *CDKN2A*/p16 knockdown in human and murine cancer cells was associated with striking Golgi compaction. *CDKN2A*/p16 knockdown sensitized cells to pharmacologic Golgi disruption via lipid peroxidation both *in vitro* and *in vivo*. Notably, reduced *Retinoblastoma*-associated protein (*RB1*) expression did not recapitulate the increased sensitivity to Golgi disruption, supporting a role for non-canonical p16 functions in this vulnerability. Together, these findings uncover a previously unrecognized link between *CDKN2A*/p16^Low^ status and Golgi remodeling, revealing a Golgi-associated metabolic vulnerability in *CDKN2A*/p16^Low^ melanomas.

## INTRODUCTION

Cyclin-dependent kinase inhibitor 2A (*CDKN2A*) is one of the most frequently altered tumor suppressor loci across human cancers, with loss or inactivation of its p16^INK4A^ product (hereafter p16) occurring in ∼40% of melanomas and ∼50% of human cancers^1, 2^. Canonically, p16 inhibits CDK4/6 to restrain RB1 phosphorylation and cell cycle progression^3^. However, accumulating evidence from our laboratory and others indicates that the tumor-suppressive functions of p16 extend beyond cell cycle control to include regulation of cellular metabolism and stress responses^4–15^. These non-canonical activities remain less well understood but raise the possibility that loss of p16 may have broader consequences for cellular physiology than previously appreciated^16, 17^. Despite the high prevalence of *CDKN2A*/p16 alterations, there are currently no therapies specifically targeting tumors with reduced or lost p16 expression, and clinical strategies largely focus on downstream cell cycle machinery^18, 19^. Identifying therapeutic strategies that exploit additional cancer cell-intrinsic consequences of *CDKN2A*/p16 loss are therefore essential for improving outcomes in this large patient population.

The Golgi apparatus (hereafter Golgi) is a highly organized perinuclear organelle that coordinates protein processing, lipid homeostasis, and vesicular trafficking^20^. Golgi morphology is closely linked to its function, and alterations in its architecture affect its processes^20^. Golgi morphology is frequently altered in cancer cells, where they can exhibit either compacted juxtanuclear morphology or fragmented, dispersed morphology^21, 22^. These morphological changes have been linked to a variety of cellular consequences, including cellular stress responses^23, 24^. It is unclear whether *CDKN2A*/p16 alters Golgi morphology and if that creates an exploitable vulnerability.

Here, we identify a previously unrecognized vulnerability of cancer cells with loss of *CDKN2A*/p16 expression (hereafter *CDKN2A*/p16^Low^) associated with altered Golgi morphology. We show that *CDKN2A*/p16^Low^ melanoma cells exhibit striking Golgi compaction, which is associated with increased sensitivity to pharmacological Golgi disruption. Notably, these phenotypes were not recapitulated in cells and tumors with decreased *RB1* gene expression, suggesting that this reflects a non-canonical function of p16 beyond the RB1–E2F pathway. Mechanistically, we demonstrate that Golgi disruption promotes lipid peroxidation–mediated cell death in *CDKN2A*/p16^Low^ cells. Consistent with these findings, pharmacological Golgi disruption selectively increases lipid peroxidation and suppresses melanoma growth *CDKN2A*/p16^Low^ tumors *in vivo*. Together, our findings identify Golgi remodeling as a therapeutically exploitable vulnerability associated with *CDKN2A*/p16 loss and support Golgi disruption as a potential precision medicine strategy for these tumors.

## RESULTS

### Golgi-associated vulnerabilities and altered morphology characterize *CDKN2A*/p16^Low^ melanoma

Beyond its canonical role in cell cycle regulation, loss or reduced expression of *CDKN2A*/p16 (hereafter *CDKN2A*/p16^Low^) has been shown to exert broader effects on cellular physiology and create selective vulnerabilities in cancer cells^4, 5, 7, 8^. Given its high prevalence across human cancers, including melanoma (∼40% of patients)^1, 2^, defining additional dependencies associated with *CDKN2A*/p16^Low^ remains a central goal. To explore such dependencies, we reanalyzed our previously published metabolic CRISPR knockout screen performed in isogenic YUMM5.2 mouse melanoma cells with *Cdkn2a* knockdown (sh*Cdkn2a*) or control (shControl)^7^ (**Fig. 1A**). Gene Set Enrichment Analysis (GSEA) of genes that significantly dropped out in sh*Cdkn2a* cells (**Table S1**) revealed a striking enrichment of Golgi and organellar membrane–associated Gene Ontology Cellular Component (GO CC) categories (**Fig. 1B** and **Table S2**). Consistent with these findings, DepMap^25^ CRISPR screening data revealed greater dependency on Golgi-associated genes in *CDKN2A*^Low^ versus *CDKN2A*^High^ cell lines across diverse tumor types with wild-type *CDKN2A* (**Fig. S1A**). Together, these findings identified Golgi membrane organization as a potential vulnerability of *CDKN2A*/p16^Low^ cells. To functionally assess the Golgi architecture in *CDKN2A*/p16^Low^ cells, we performed immunofluorescence staining for GM130, a well-established cis-Golgi membrane marker^26, 27^. Knockdown of *Cdkn2a* (shRNAs targeting *Cdkn2a* exon 3) in YUMM5.2 murine melanoma cells (**Fig. S1B**) and of p16 (shRNAs targeting *CDKN2A* exon 1α) in SK-MEL-28 human melanoma cells (**Fig. S1C**) was associated with a marked reorganization of the Golgi, shifting from a perinuclear morphology to a compacted, highly condensed juxtanuclear structure (**Fig. 1C-D**). *Cdkn2a*/p16 knockdown did not alter total GM130 protein abundance (**Fig. S1B-C**) and was not accompanied by broad changes in the expression of Golgi-related genes in either model (**Fig. S1D-E and Table S3-4**) in either model, suggesting that the observed Golgi phenotype is not associated with changes in total Golgi mass or transcriptional regulation of Golgi components downstream of *CDKN2A* loss. Together, these data indicate that altered Golgi morphology is associated with *CDKN2A*/p16^Low^ status and support Golgi integrity as a potential vulnerability in this disease context.

**Figure 1.**
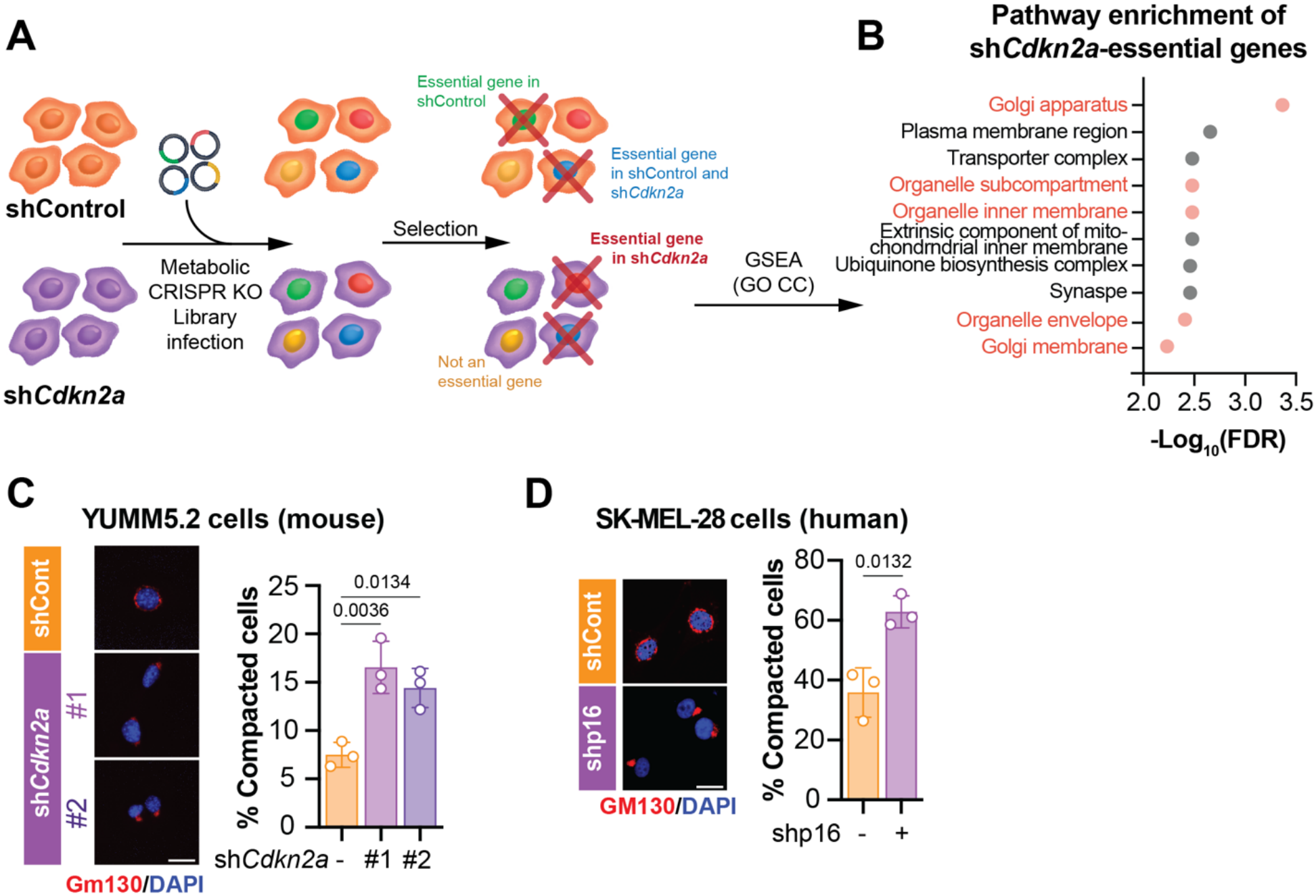
CRISPR dropout screen uncovers the Golgi as a potential vulnerability in *Cdkn2a*/p16^Low^ cancer cells, which is associated with compacted Golgi morphology. **(A)** Schematic overview of the metabolically-focused CRISPR knockout (KO) screen performed in YUMM5.2 mouse melanoma cells with knockdown of *Cdkn2a* (sh*Cdkn2a*) or control (shControl). **(B)** Gene Set Enrichment Analysis (GSEA) of Gene Ontology Cellular Component (GOCC) pathways using genes significantly depleted in sh*Cdkn2a* cells relative to shControl cells in the CRISPR KO screen (defined as genes targeted by >5 sgRNAs with p<0.05). Golgi and other organelle-related signatures are depicted in red. **(C-D)** Representative immuno-fluorescence images (left) and quantification (right) of the Golgi marker GM130 in **(C)** YUMM5.2 cells with knockdown of *Cdkn2a* (sh*Cdkn2a,* hairpins #1 and #2) or control (shControl), and **(D)** SK-MEL-28 human melanoma cells with knockdown of p16 (shp16) or control (shControl). Nuclei are counterstained with DAPI. Mean ± SD; one-way ANOVA followed by Šídák post-hoc test and student’s t-test. Data shown are representative of three independent experiments. Scale bars = 50μm.

### Perturbation of compacted Golgi morphology is a vulnerability of *CDKN2A*/p16^Low^ melanoma cells

We found that *CDKN2A*/p16^Low^ melanoma cells exhibit a compacted, juxtanuclear Golgi morphology that differs from the perinuclear organization observed in *CDKN2A*/p16^High^ cells (**Fig. 1C-D**). Prior studies have shown that deviations from normal Golgi architecture reduce resilience to Golgi-specific stress, increasing sensitivity to pharmacologic Golgi disruption^28–32^. We therefore reasoned that the compacted Golgi morphology in *CDKN2A*/p16^Low^ melanoma cells may represent an exploitable structural vulnerability. To test this hypothesis, we perturbed Golgi architecture using three mechanistically distinct Golgi-disrupting agents^32^: Brefeldin A (BFA) and Golgicide A (GCA), which inhibit ARF1-dependent vesicle trafficking, and monensin (Mon), a Na⁺/H⁺ ionophore that disrupts Golgi function by collapsing ionic gradients and pH (**Fig. 2A**). Immunofluorescence analysis of GM130 revealed that sh*Cdkn2a*/shp16 cells were substantially more susceptible to Golgi disruption than isogenic controls (**Fig. 2B-C** and **Fig. S2A-B**). This increased sensitivity was observed across both murine YUMM5.2 and human SK-MEL-28 melanoma models and with all three compounds, indicating that the phenotype is not restricted to a particular cell model or Golgi-disrupting agent.

**Figure 2.**
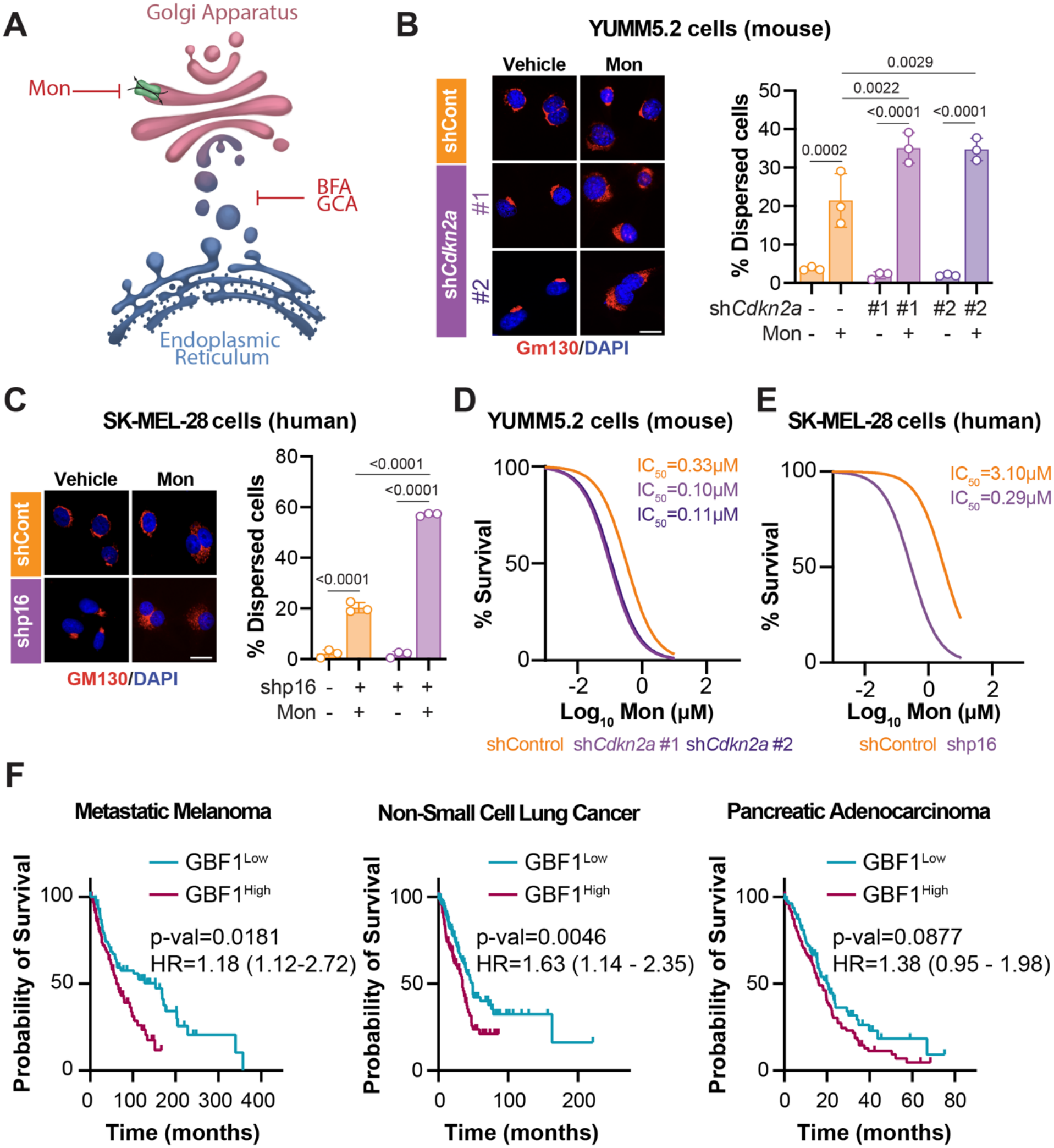
Golgi-disrupting agents enhance Golgi dispersion and reduce viability in *Cdkn2a*/p16^Low^ cancer cells. **(A)** Schematic illustrating the mechanism of action of the Golgi-disrupting agents used in this study: Brefeldin A (BFA), Golgicide A (GCA), and Monensin (Mon). **(B-C)** Representative immunofluorescence images (left) and quantification (right) of the Golgi marker GM130 following treatment with Monensin (Mon) in **(B)** YUMM5.2 cells with knockdown of *Cdkn2a* (sh*Cdkn2a,* hairpins #1 and #2) or control (shCont), and **(D)** SK-MEL-28 cells with knockdown of p16 (shp16) or control (shCont). Nuclei are counterstained with DAPI. Mean ± SD; one-way ANOVA followed by Šídák post-hoc test. Data shown are representative of three independent experiments. Scale bars = 25 µm. **(D-E)** Monensin (Mon) dose-response curve and IC_50_ value. **(D)** YUMM5.2 cells with knockdown of *Cdkn2a* (sh*Cdkn2a,* hairpins #1 and #2) or control (shControl), and **(E)** SK-MEL-28 cells with knockdown of p16 (shp16) or control (shControl). All data shown are representative of three independent experiments. **(F)** Overall survival of the indicated cancer patients from TCGA stratified based on *GBF1* median expression level. Log-rank test. HR = hazard ratio.

We next asked whether this structural vulnerability could be pharmacologically exploited to selectively impair the viability of *CDKN2A*/p16^Low^ melanoma cells. sh*Cdkn2a*/shp16 cells showed significantly lower IC_50_ values for Golgi-disrupting agents than control cells in both murine and human melanoma models (**Fig. 2D-E** and **Fig. S2C-D**). Consistent with these findings, *CDKN2A*^Low^ cell lines from the DepMap^25^ showed modest but significantly increased sensitivity to Golgi disruptors compared with *CDKN2A*^High^ cells (**Fig. S2E**). These findings support the idea that the compacted Golgi observed in *Cdkn2a*/p16^Low^ cells represents a structural vulnerability that can be therapeutically exploited. We and others have established that *CDKN2A*/p16 has functions beyond its canonical role in the RB1–E2F pathway^7, 8, 16^. We therefore asked whether the sensitivity to Golgi disruption in *CDKN2A*/p16^Low^ cells results from loss of canonical p16–RB1–E2F signaling or from a non-canonical function of *CDKN2A*/p16. If this vulnerability were mediated through the canonical p16–RB1–E2F pathway, loss of *RB1* would phenocopy *CDKN2A*/p16 loss. We therefore compared Golgi-disruptor sensitivity in *RB1*^Low^ and *RB1*^High^ cell lines from the DepMap^25^. Unlike *CDKN2A*^Low^ cells, *RB1*^Low^ cells were not more sensitive to Golgi disruption (**Fig. S2F**), suggesting that this vulnerability is not a consequence of impaired canonical p16–RB1–E2F signaling but rather reflects a non-canonical function of *CDKN2A*/p16.

Finally, we asked whether Golgi dispersion is associated with clinical outcomes, independent of *CDKN2A*/p16 status. We assessed expression of the cis-Golgi ARF guanine nucleotide exchange factor GBF1 as a proxy for Golgi dispersion. GBF1 is required for COPI vesicle formation and maintenance of Golgi architecture, and its inhibition results in rapid Golgi dispersion and functional colapse^33^ (**Fig. 2A**). Analysis of TCGA datasets revealed that low *GBF1* expression (*GBF1*^Low^) was associated with significantly better overall survival in melanoma, pancreatic cancer, and non-small cell lung cancer (**Fig. 2F**). Together, these findings suggest that Golgi morphology represents a therapeutic vulnerability beyond *CDKN2A*/p16^Low^ melanoma.

### Golgi disruption primes *CDKN2A*/p16^Low^ melanoma cells for lipid peroxidation–mediated cell death

We next sought to determine the mechanism underlying the heightened sensitivity of *CDKN2A*/p16^Low^ cancer cells to Golgi disruption. We performed RNA-Seq on isogenic murine YUMM5.2 and human SK-MEL-28 melanoma cells treated with monensin or DMSO. Surprisingly, only 2 differentially expressed genes overlapped between the two models, with no clear enrichment for a common biological pathway (**Fig. S3A** and **Table S5**). To determine whether this limited overlap reflected differences between species or cell types, we extended our analysis to publicly available RNA-Seq data from monensin-treated THP-1 human acute monocytic leukemia cells (GSE130657)^34^ and A549 human lung carcinoma cells^35^, both of which harbor homozygous *CDKN2A* deletions^36, 37^. Strikingly, no differentially expressed genes were shared across all four cell lines following monensin treatment (**Fig. 3B** and **Table S6**). These findings indicate that monensin does not elicit a common transcriptional response across these models suggesting that the selective sensitivity of *CDKN2A*/p16^Low^ cells to Golgi disruption is unlikely to be explained by a shared alteration of the transcriptional program. We therefore explored alternative mechanisms.

**Figure 3.**
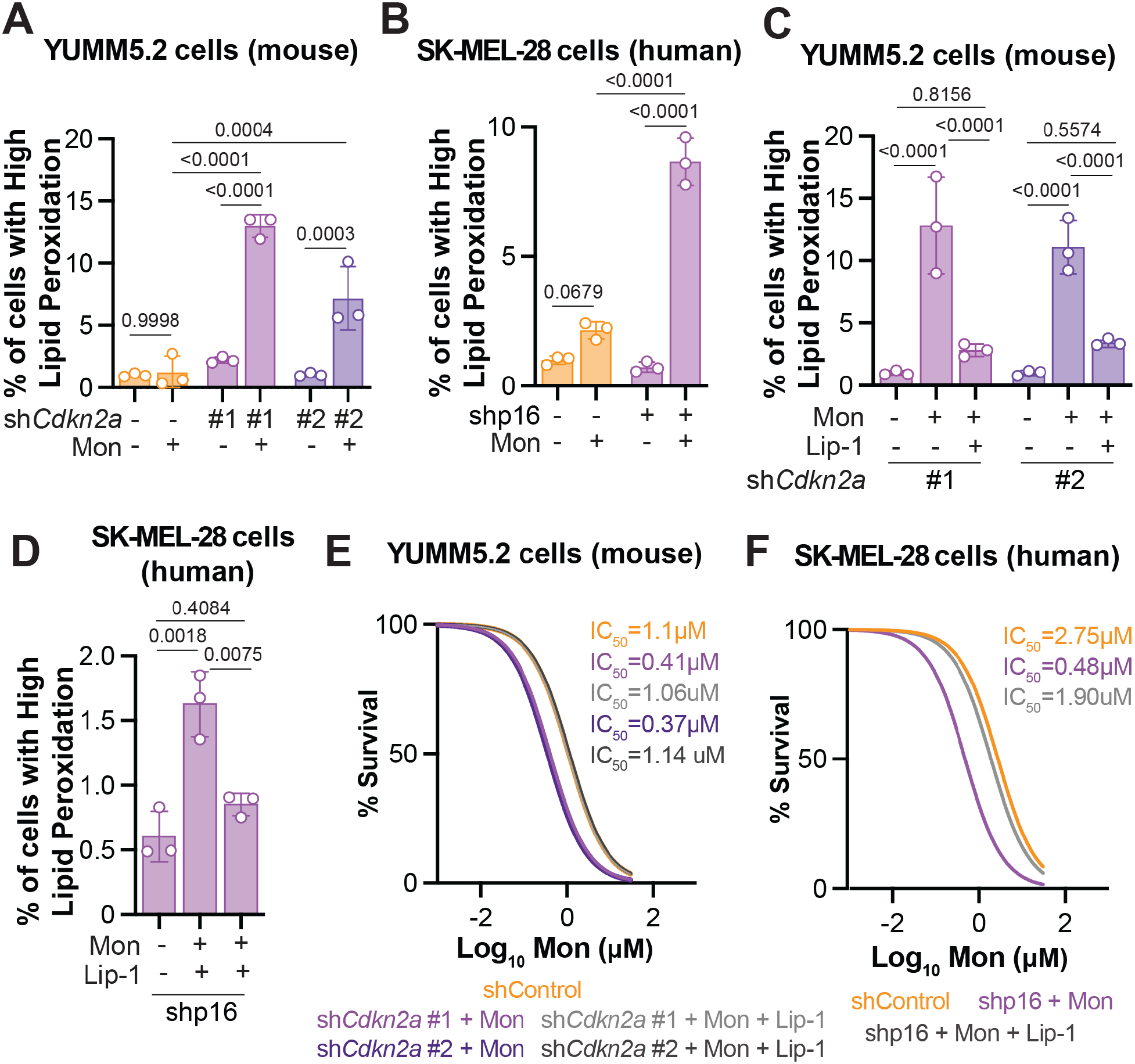
Golgi disruption induces lipid peroxidation-mediated cell death in *Cdkn2a*/p16^Low^ melanoma cells. **(A-B)** Quantification of lipid peroxidation in **(A)** YUMM5.2 cells expressing shControl or sh*Cdkn2a* (hairpins #1 and #2) and **(B)** SK-MEL-28 cells expressing shControl or shp16 following treatment with Monensin (Mon). Lipid peroxidation was assessed by C11-BODIPY581/591 staining followed by flow cytometry. Mean ± SD; one-way ANOVA followed by Šídák post-hoc test. Data shown are representative of three independent experiments. **(C-D)** Quantification of lipid peroxidation in **(C)** YUMM5.2 sh*Cdkn2a* (hairpins #1 and #2) cells and **(D)** SK-MEL-28 shp16 cells treated with Mon alone or in combination with Lip-1. Lipid peroxidation was assessed by C11-BODIPY581/591 staining followed by flow cytometry. Mean ± SD; one-way ANOVA followed by Šídák post-hoc test. Data shown are representative of three independent experiments. **(E-F)** Monensin dose-response curves and IC_50_ values for Mon alone or in combination with Lip-1 in **(E)** YUMM5.2 cells with knockdown of *Cdkn2a* (sh*Cdkn2a*, hairpins #1 and #2) and **(F)** SK-MEL-28 cells with knockdown of p16 (shp16). Data shown are representative of three independent experiments.

Prior studies have shown that Golgi-disrupting agents induce lipid peroxidation^28, 38^. Based on these observations, we hypothesized that Golgi disruption in *CDKN2A*/p16^Low^ cells promotes lipid peroxidation–mediated cell death. To test this hypothesis, shControl and sh*Cdkn2a*/shp16 cells were treated with Golgi-disrupting agents at concentrations that induce differential Golgi dispersion (**Fig. 2B-C**), and lipid peroxidation was assessed using the C11-BODIPY^581/591^ fluorescent probe^39^. Across all three Golgi disruptors, sh*Cdkn2a*/shp16 cells exhibited significantly higher lipid peroxidation than isogenic controls in both murine YUMM5.2 and human SK-MEL-28 models (**Fig. 3C-D** and **Fig. S4A-B**). Interestingly, monensin induced broadly similar changes in the expression of major antioxidant and ferroptosis-associated genes in sh*Cdkn2a*/shp16 and shControl cells (**Fig. S4C-D** and **Table S7-8**). These findings suggest that the increased sensitivity of *CDKN2A*/p16^Low^ cells to Golgi disruption is not simply driven by differential transcriptional regulation of these pathways downstream of loss of *CDKN2A*/p16.

Next, we tested whether the increased BODIPY581/591 signal reflected *bona fide* lipid peroxidation by treating cells with the lipid peroxidation inhibitor liproxstatin-1 (Lip-1)^40^. Lip-1 abrogated the increase in BODIPY581/591 staining induced by Golgi disruption (**Fig. 3D**) and restored the IC_50_ values of sh*Cdkn2a*/shp16 cells to control levels (**Fig. 3E-F**). These findings demonstrate that lipid peroxidation is a primary driver of the increased sensitivity of *CDKN2A*/p16^Low^ melanoma cells to Golgi disruption. Finally, we examined the directionality of the relationship between Golgi disruption and lipid peroxidation. Lip-1 did not reverse monensin-induced Golgi dispersion (**Fig. S4E-F**). Conversely, pharmacologic induction of lipid peroxidation using the GPX4 inhibitor RSL3^40^ failed to induce Golgi dispersion (**Fig. S4G-H**). Altogether, these findings indicate that Golgi disruption primes *CDKN2A*/p16^Low^ cells for lipid peroxidation–mediated death, which is not associated with altered ferroptosis and antioxidant expression programs.

### Golgi disruption decreases *Cdkn2a*/p16^Low^ tumor growth *in vivo* and is associated with lipid peroxidation

To determine whether the vulnerability of *CDKN2A*/p16^Low^ cells to Golgi disruption extends *in vivo*, we generated subcutaneous tumors using isogenic YUMM5.2 melanoma cells in immunocompetent mice and SK-MEL-28 cells in immunocompromised mice. Mice were then treated with the Golgi disruptor monensin (**Fig. 4A**), which has previously been used *in vivo*^29^. Monensin treatment significantly reduced tumor growth in sh*Cdkn2a*/p16 tumors, whereas shControl tumors were largely unaffected (**Fig. 4B** and **Fig. S5A**). This selective effect was not accompanied by significant changes in mouse body weight (**Fig. S5B-C**), suggesting that the reduced tumor growth was not due to overt systemic toxicity. Consistent with our *in vitro* findings, monensin-treated sh*Cdkn2a* tumors also exhibited significantly higher 4-hydroxynonenal (4-HNE) staining than monensin-treated shControl tumors (**Fig. 4C**), indicating increased lipid peroxidation *in vivo*. Together, these findings demonstrate that Golgi disruption selectively suppresses *CDKN2A*/p16^Low^ tumor growth and is associated with increased lipid peroxidation *in vivo*. Collectively, our data support a model in which *CDKN2A*/p16 loss creates a Golgi-dependent vulnerability whereby Golgi disruption promotes lipid peroxidation and constrains tumor growth (**Fig. 4D**).

**Figure 4.**
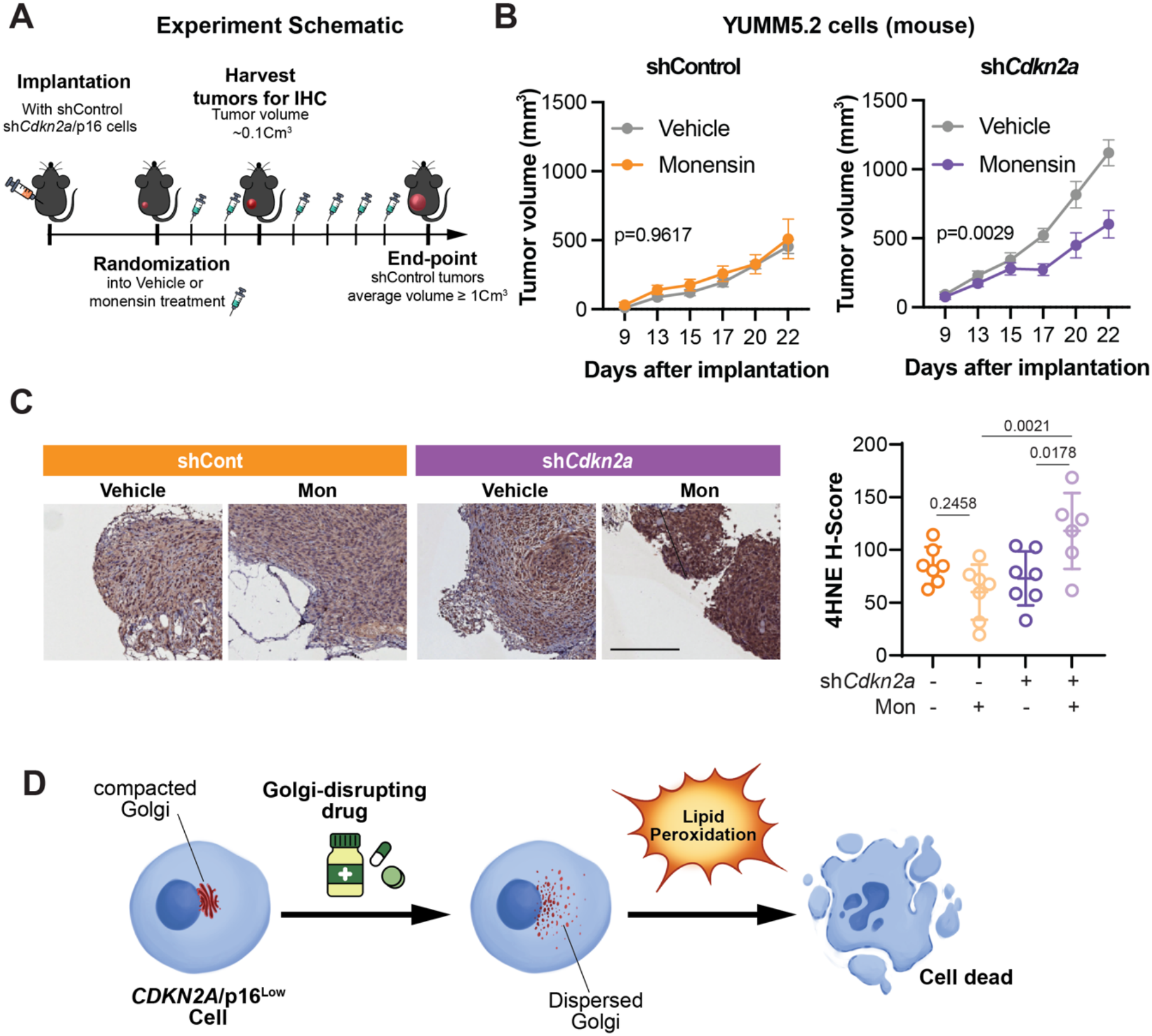
Golgi disruption promotes lipid peroxidation and suppresses tumor growth *in vivo* and is associated with favorable outcomes in *CDKN2A*/p16^Low^ tumors. **(A)** Schematic overview of the *in vivo* experiment design. **(B-C)** YUMM5.2 shControl and sh*Cdkn2a* cells were implanted in immunocompetent mice. **(B)** Tumor growth curves of tumors treated treated with vehicle or Monensin (10mg/kg, i.p, 3x/week for a total of 6 doses). Mixed-effects model (day x group – Mon Vs. DMSO) in the indicated comparison. **(C)** Representative immunohistochemical images (left) and quantification (right) of 4-hydroxynonenal (4HNE) staining in YUMM5.2 shControl and sh*Cdkn2a* tumors treated with vehicle or Monensin (10mg/kg, i.p, 2 total doses). Mean ± SD; one-way ANOVA followed by Šídák post-hoc test. **(D)** Proposed model illustrating how *CDKN2A*/p16 loss induces a compacted Golgi architecture that creates a vulnerability to pharmacological Golgi disruption, resulting in increased lipid peroxidation and cell death.

## DISCUSSION

The role of *CDKN2A*/p16 as a critical regulator of the cell cycle has been well established for more than three decades^3, 41^. However, accumulating evidence from our laboratory and others indicates that *CDKN2A*/p16 also exerts non-canonical functions that extend beyond cell cycle control, including regulation of cellular metabolism^4, 7, 8^ and organelle function, such as mitochondrial homeostasis^10, 11^. Here, we identify a previously unrecognized link between *CDKN2A*/p16 loss and Golgi architecture, demonstrating that *CDKN2A* /p16^Low^ cancer cells adopt a compacted Golgi morphology (**Fig. 1**) that confers heightened sensitivity to pharmacologic Golgi disruption (**Fig. 2**), leading to lipid peroxidation-mediated cell death *in vitro* (**Fig. 3**) and reduced tumor growth *in vivo* (**Fig. 4**). Collectively, these findings suggest that targeting altered Golgi architecture may represent a therapeutic opportunity for the substantial fraction (∼50%) of human cancers characterized by *CDKN2A*/p16 loss, including the ∼40% of melanomas^1^.

Despite the high prevalence of *CDKN2A*/p16 loss across human cancers, there are currently no therapies specifically designed to target *CDKN2A*/p16^Low^ tumors^18, 19^. Clinical management instead relies largely on downstream inhibition of cell cycle regulators or treatment strategies that are agnostic to p16 status^42^, leaving the non-canonical consequences of p16 loss largely therapeutically unexplored. This limitation is further compounded by the infrequent assessment of *CDKN2A*/p16 status in routine clinical practice, particularly in melanoma outside the context of suspected hereditary disease^43^. In this setting, the present findings provide proof-of-concept that *CDKN2A*/p16^Low^ tumors harbor therapeutically exploitable, cancer cell-intrinsic liabilities that extend beyond cell cycle dysregulation. While monensin is not approved for human use, it is FDA-approved for veterinary applications as an ionophore antibiotic and has demonstrated a favorable safety profile over decades of use in livestock, including cattle and poultry^38, 44–46^. Its prior efficacy in preclinical cancer models *in vivo*^32, 47, 48^, together with the selective growth suppression of *CDKN2A*/p16^Low^ tumors observed here (**Fig. 4**), supports the concept that Golgi disruption represents a tractable therapeutic strategy for this genetically defined tumor subset. Importantly, this vulnerability is not limited to a single compound: *CDKN2A*/p16^Low^ cancer cells display increased sensitivity to multiple mechanistically distinct Golgi-disrupting agents, and analysis of DepMap datasets further supports a broader dependency on Golgi integrity across diverse *CDKN2A*^Low^ cancer cell lines (**Fig. S2E**), arguing against a compound-specific effect. Consistent with these experimental findings, analysis of patient datasets revealed that low expression of the cis-Golgi ARF guanine nucleotide exchange factor *GBF1*—a key regulator of Golgi structural maintenance^33^—is associated with improved overall survival across multiple cancer types, including melanoma (**Fig. 2F**). Importantly, our findings suggest that the increased sensitivity to Golgi disruption observed in *CDKN2A*/p16^Low^ cells reflects a non-canonical function of p16, as this sensitivity was not recapitulated by loss of *RB* (**Fig. S2F**). Together, these data reinforce the clinical relevance of Golgi integrity and suggest that tumors with reduced Golgi maintenance capacity are intrinsically more vulnerable to stress-inducing perturbations, positioning Golgi disruption as a generalizable, cancer cell–intrinsic vulnerability in *CDKN2A*/p16^Low^ tumors. Importantly, the efficacy of Golgi disruption in both immunocompetent and immunocompromised models suggest that this vulnerability is at least in part independent of immune context and reflects intrinsic stress sensitivity of *Cdkn2a*/p16^Low^ cancer cells (**Fig. 4 and Fig. S5A**). This observation is particularly relevant given accumulating evidence that *CDKN2A*/p16^Low^ tumors respond poorly to immune checkpoint blockade^49–52^, despite the fact that immunotherapy has become standard of care for multiple *CDKN2A*/p16^Low^-associated cancers, including melanoma and non-small cell lung cancer (NSCLC)^53, 54^. Our findings also raise the possibility that Golgi disruption strategies could be combined with immunotherapy to enhance therapeutic efficacy. In support of this concept, our laboratory has previously shown that enhancing anti-PD-1-mediated macrophage phagocytosis limits tumor growth in *CDKN2A*^Low^ cancers^5^. Together, these findings suggest that combining macrophage-mediated tumor clearance with cancer cell-intrinsic killing mechanisms, such as Golgi disruption-induced lipid peroxidation, may provide a complementary specific therapeutic strategy for *CDKN2A*/p16^Low^ tumors.

The Golgi apparatus is increasingly recognized as a dynamic organelle whose structural organization is tightly coupled to lipid handling, vesicular trafficking, and stress adaptation^20^. Here, we show that *CDKN2A*/p16^Low^ cancer cells adopt a compacted Golgi morphology (**Fig. 1C-D**) that is selectively vulnerable to pharmacologic disruption (**Fig. 2D-E**), leading to elevated lipid peroxidation and cell death (**Fig. 3**). Emerging evidence has linked Golgi stress to lipid peroxidation-dependent cytotoxicity^28^. Prior literature has shown that oxidative stress can induce Golgi fragmentation through different mechanisms including cytoskeletal damage or ER stress^55^. However, our data demonstrate that inhibition of lipid peroxidation rescues cell viability without restoring Golgi structure (**Fig. 3E-F** and **S4E-F**), supporting a model in which Golgi disruption acts upstream to promote oxidative lipid damage that ultimately mediates cell death. These observations are consistent with emerging evidence linking Golgi stress to lipid peroxidation-dependent cytotoxicity^28^. Thus, it is likely that both directions can occur in a context-dependent manner, and further research is needed to define the upstream oncogenic determinants that dictate the directionality of this relationship.

In contrast to a recent study showing that *CDKN2A* loss is sufficient to increase lipid peroxidation and ferroptosis sensitivity in glioblastoma^56^, we did not observe consistent differences in basal lipid peroxidation between sh*Cdkn2a*/p16 and shControl cells (**Fig. 3A and Fig. S4A-B**). Our data support a model in which Golgi disruption acts upstream to promote lipid peroxidation-mediated death in *CDKN2A*/p16^Low^ cells (**Fig. S4E-H**). However, the mechanisms linking Golgi structural perturbation to lipid peroxidation remain to be defined in our system. Given the central role of the Golgi in lipid synthesis and trafficking^20^, it is plausible that Golgi disruption alters cellular lipid homeostasis thereby increasing the availability of oxidation-prone membrane lipids (e.g. polyunsaturated fatty acids)^56^. It is also possible that Golgi fragmentation disrupts normal vesicular trafficking of the iron machinery promoting the sensitization of *CDKN2A*/p16^Low^ cells to oxidative lipid damage once Golgi integrity is pharmacologically challenged. Defining how the altered Golgi state established by *CDKN2A*/p16 loss affects lipid composition, antioxidant capacity and redox homeostasis will be an important direction for future studies.

Beyond its role in cell survival, Golgi organization is central to the secretory capacity of cancer cells^57^. Interestingly, our laboratory has previously shown that loss of p16 reprograms the secretion of cytokines and chemokines with established roles in regulating the tumor microenvironment^6^. An intriguing possibility raised by the present work is that Golgi compaction in p16^Low^ cells contributes to this secretory reprogramming. While our *in vivo* data using a xenograft human cell line model suggest that Golgi disruption-mediated tumor growth inhibition occurs at least in part independently of the immune compartment, these experiments do not exclude contributions from non-immune components of the tumor microenvironment, such as cancer-associated fibroblasts, innervation, or other stromal elements whose behavior can be modulated by cancer cell-derived secreted factors. Thus, defining how altered Golgi architecture in *CDKN2A*/p16^Low^ cells integrates cell-intrinsic secretory programs with tumor-stromal communication represents an important and exciting direction for future investigation.

In summary, this study links loss of the tumor suppressor *CDKN2A*/p16 to a compacted Golgi state that creates a selective therapeutic vulnerability to lipid peroxidation due to Golgi dispersion. The selective impairment of *Cdkn2a*/p16^Low^ tumor growth *in vivo* highlights Golgi integrity as a tractable therapeutic target in this large subset of cancers (∼40% of melanoma and ∼50% of all human cancer) that currently lack tailored treatment strategies. Beyond *CDKN2A*/p16 loss, this work reinforces emerging evidence that organelle-specific states can be exploited as a new axis for precision oncology, positioning Golgi architecture as a functional node connecting genetic lesions, oxidative stress, and therapeutic response.

## SUPPLEMENTAL FIGURES

**Figure S1.**
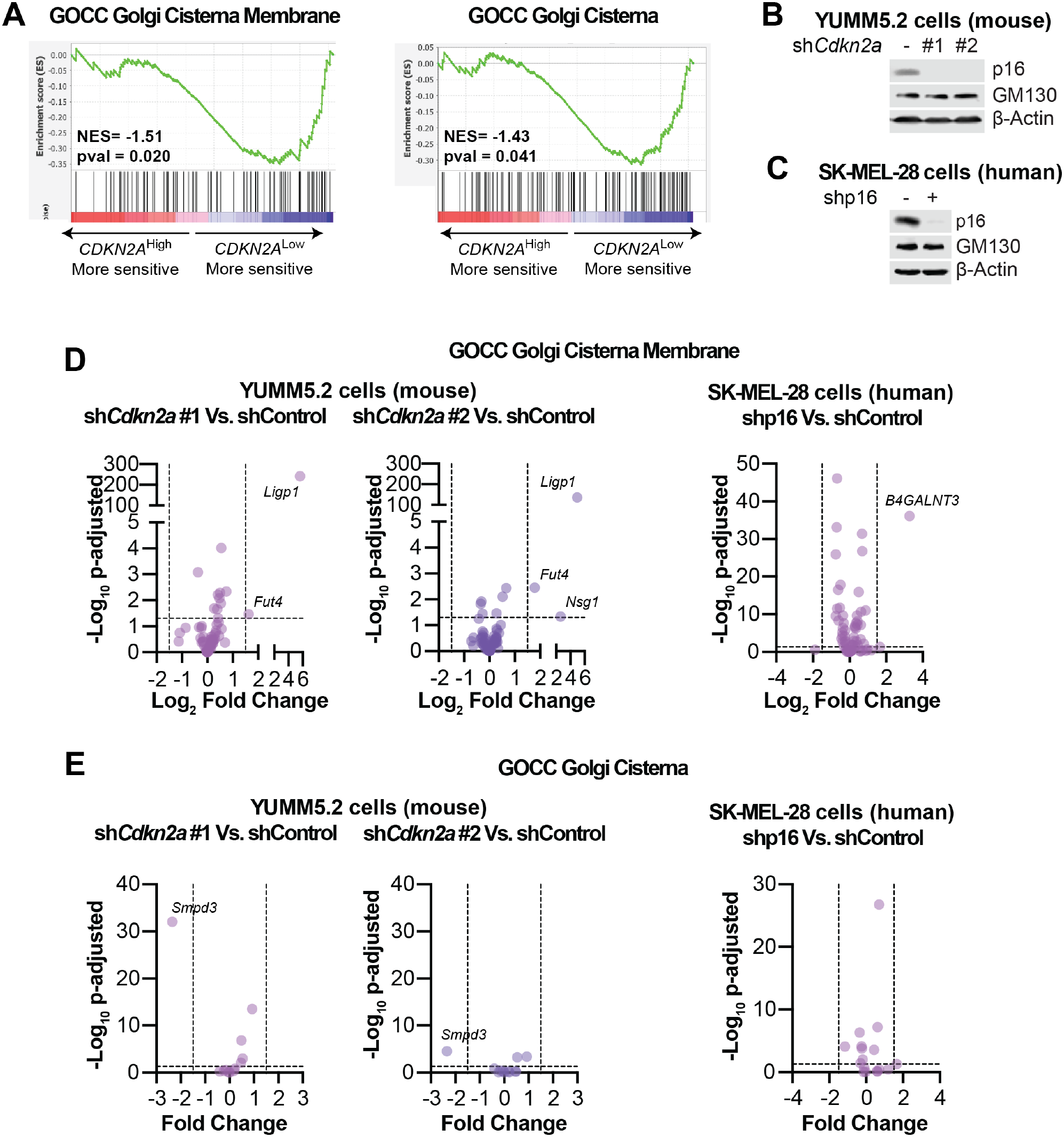
*CDKN2A*^Low^ cancer cells are more sensitive to knockout of Golgi-associated genes and knockdown of *Cdkn2a* or p16 does not alter total GM130 protein abundance. Related to Figure 1. **(A)** Analysis of DepMap CRISPR data demonstrate that *CDKN2A*^Low^ cancer cells are more sensitive to knockout of Golgi-associated genes that *CDKN2A*^High^ cells. **(B-C)** Immunoblot analysis of the indicated proteins in **(B)** YUMM5.2 cells with knockdown of *Cdkn2a* (sh*Cdkn2a*, hairpins #1 and #2) or control (shControl), and **(C)** SK-MEL-28 cells with knockdown of p16 (shp16) or control (shControl). β-Actin was used as a loading control. **(D-E)** Volcano plots of RNA-seq analysis for YUMM5.2 cells expressing shControl or sh*Cdkn2a* (hairpins #1 and #2) and SK-MEL-28 cells expressing shControl or shp16 following Monensin treatment. Shown are the expression level of genes associated to the pathways **(D)** Golgi Cisterna Membrane (GO CC), and **(E)** Golgi Cis Cisterna (GO CC).

**Figure S2.**
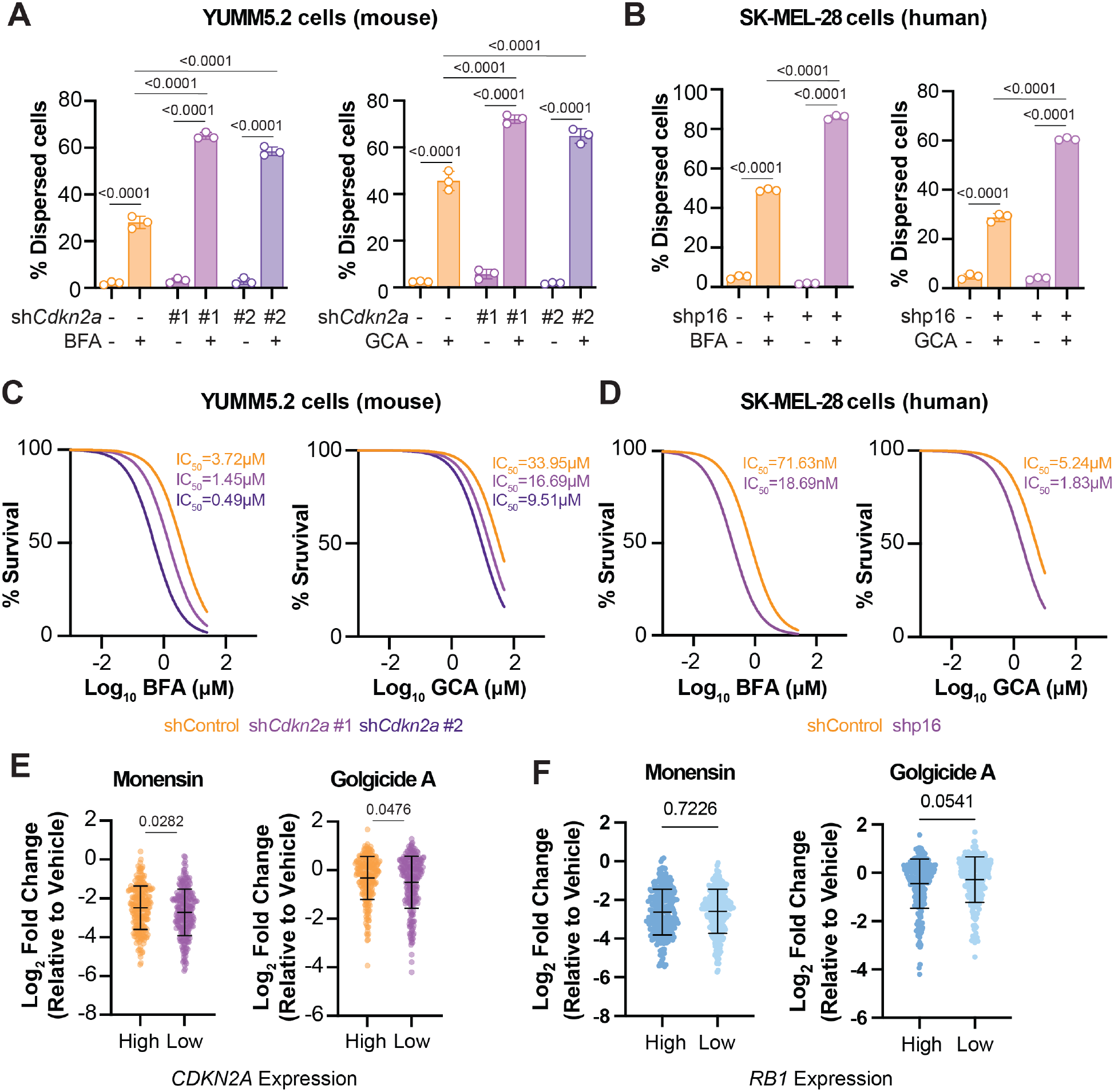
*CDKN2A*/p16^Low^ cancer cells show enhanced Golgi dispersion and increased sensitivity to Golgi disruptors. Related to Figure 2. **(A-B)** Quantification of Golgi dispersion, assessed by GM130 immunofluorescence following treatment with Brefeldin A (BFA; left) or Golgicide A (GCA; right) in **(A)** YUMM5.2 cells with knockdown of *Cdkn2a* (sh*Cdkn2a*, hairpins #1 and #2) or control (shControl), and **(B)** SK-MEL-28 cells with knockdown of p16 (shp16) or control (shControl). Mean ± SD; one-way ANOVA. Data shown are representative of three independent experiments. **(C-D)** BFA (left) and GCA (right) dose–response curves and IC_50_ values in **(C)** YUMM5.2 cells with knockdown of *Cdkn2a* (sh*Cdkn2a*, hairpins #1 and #2) or control (shControl), and **(D)** SK-MEL-28 cells with knockdown of p16 (shp16) or control (shControl). Data shown are representative of three independent experiments. **(E-F)** Dot plot showing the log_2_ fold-change in cell survival following treatment with indicated Golgi-disrupting agents across a panel of cancer cell lines stratified by (E) *CDKN2A* expression (*CDKN2A*^Low^ vs. *CDKN2A*^High^) and (F) *RB1* expression (*RB1*^Low^ vs. *RB1*^High^). Data were obtained from the DepMap database. Mean ± SD; Student’s t-test.

**Figure S3.**
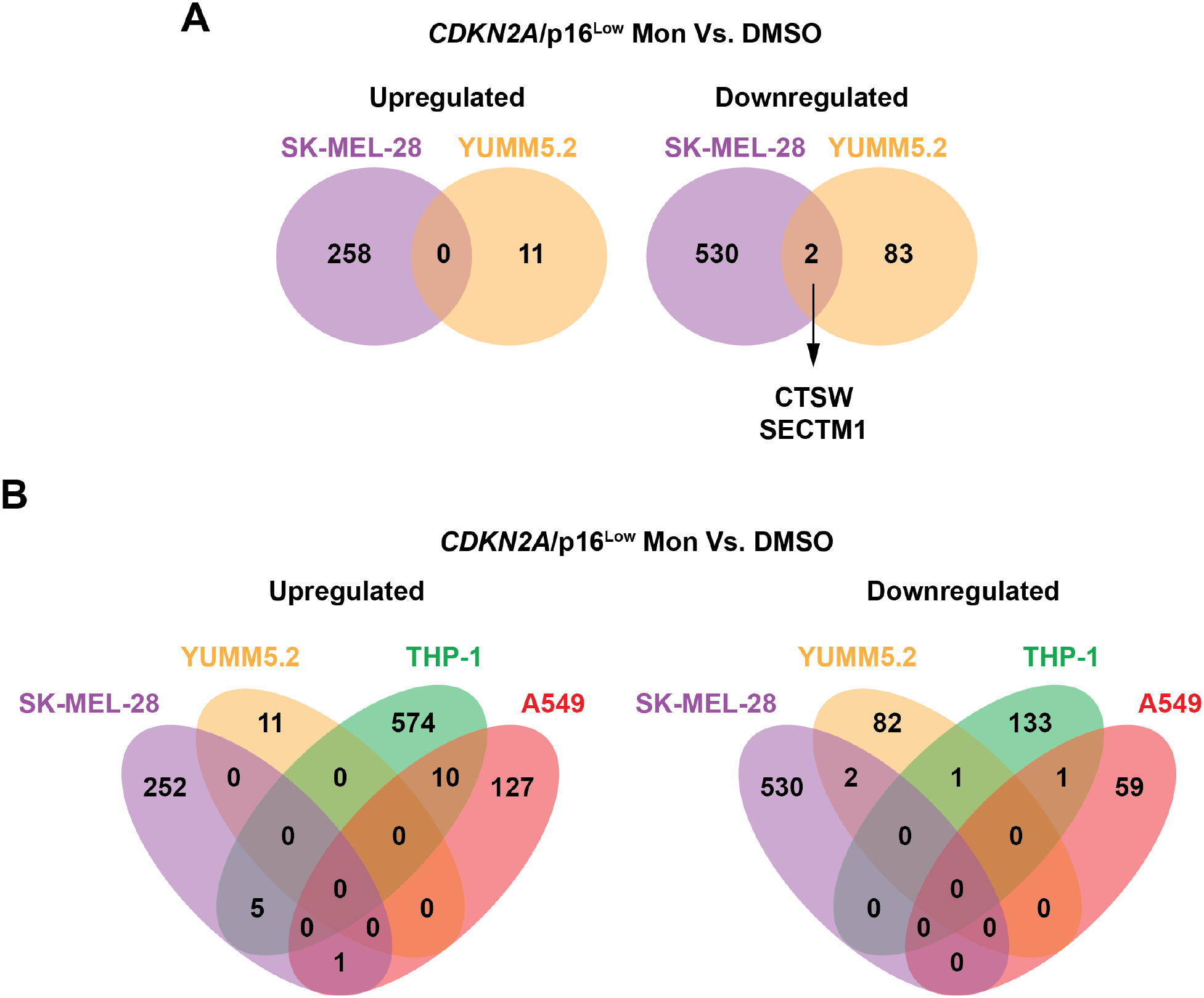
Monensin does not induce a common transcriptional response across *CDKN2A*/p16^Low^ cancer cell models. Related to Figure 3. **(A)** Venn diagrams showing the overlap of upregulated and downregulated genes following monensin (Mon) treatment in murine melanoma YUMM5.2 sh*Cdkn2a* cells and human melanoma SK-MEL-28 shp16 cells. **(B)** Venn diagrams showing the overlap of upregulated and downregulated genes following monensin treatment in the murine cell line YUMM5.2, and the human cell lines SK-MEL-28, THP-1 and A549. Numbers indicate differentially expressed genes in each overlapping or non-overlapping group. Differentially expressed genes were defined as those with an p-adj < 0.05 and a log₂ fold change ≥ 1.5 for upregulated genes or ≤ −1.5 for downregulated genes.

**Figure S4.**
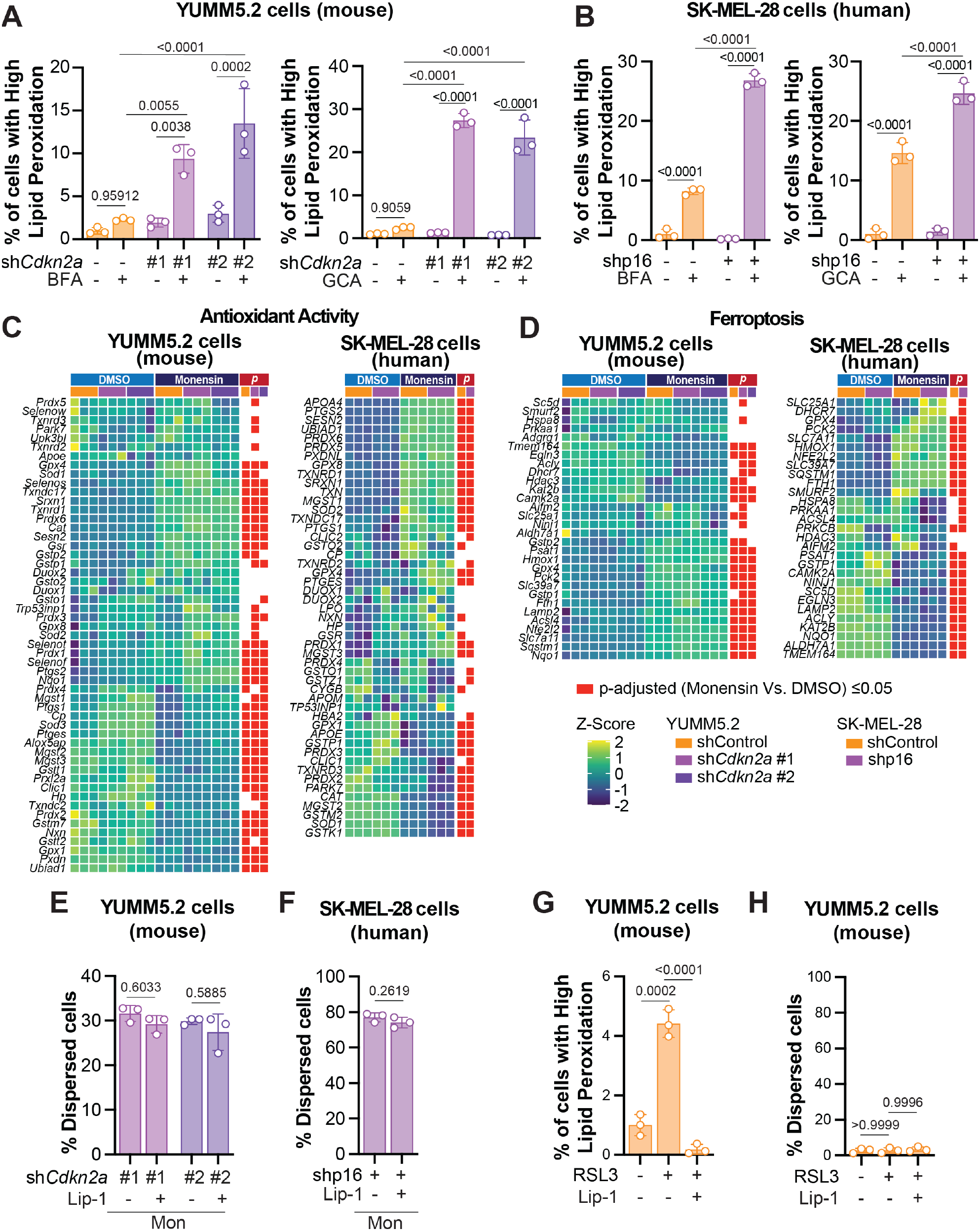
Golgi disruption induces lipid peroxidation, whereas lipid peroxidation does not induce Golgi disruption. Related to Figure 3. **(A–B)** Quantification of lipid peroxidation in **(A)** YUMM5.2 cells expressing shControl or sh*Cdkn2a* (hairpins #1 and #2) and **(B)** SK-MEL-28 cells expressing shControl or shp16 following treatment with Brefeldin A (BFA; left) and Golgicide A (GCA; right). Lipid peroxidation was assessed by C11-BODIPY581/591 staining followed by flow cytometry. Mean ± SD; one-way ANOVA. Data shown are representative of three independent experiments. **(C-D)** Heatmaps of RNA-seq analysis showing the expression of genes associated with **(C)** the Ferroptosis (Wikipathways) and **(D)** Antioxidant Activity (GO MF) gene sets in YUMM5.2 cells expressing shControl or sh*Cdkn2a* (hairpins #1 and #2) and SKMEL-28 cells expressing shControl or shp16 following monensin treatment. **(E-F)** Quantification of Golgi dispersion following treatment with Monensin alone or in combination with Lip-1 in **(E)** YUMM5.2 cells expressing sh*Cdkn2a* (hairpins #1 and #2), and **(F)** SK-MEL-28 cells expressing shp16. Mean ± SD; one-way ANOVA or student’s t-test. Data shown are representative of three independent experiments. **(G-H)** Quantification of **(G)** lipid peroxidation and **(H)** Golgi dispersion in YUMM5.2 cells expressing shControl following pharmacologic induction of lipid peroxidation with RSL3 alone or in combination with Lip-1. Mean ± SD; one-way ANOVA or student’s t-test. Data shown are representative of three independent experiments.

**Figure S5.**
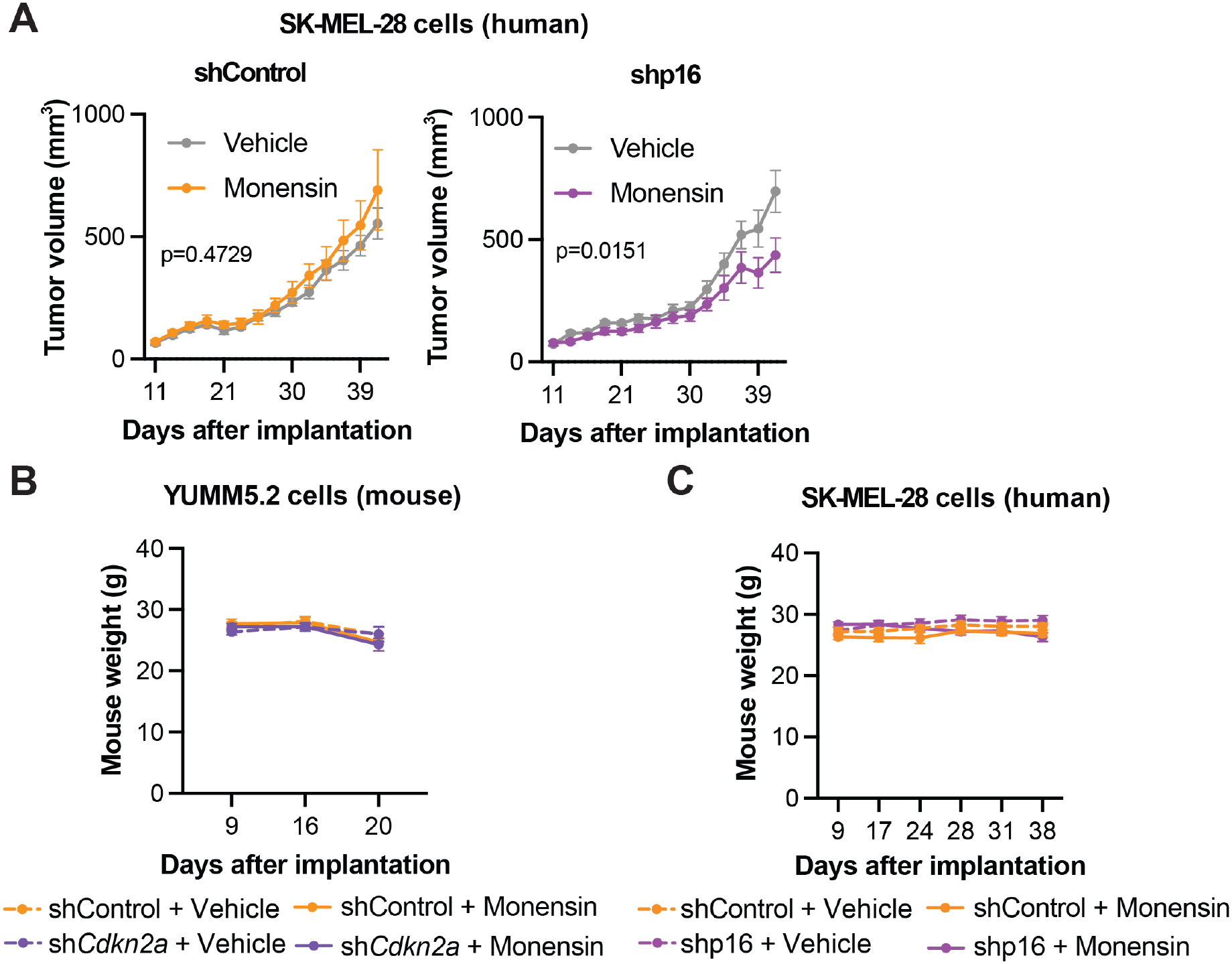
Golgi disruption suppresses tumor growth and is well tolerated *in vivo.* Related to Fig 4. **(A)** Tumor growth curves following monensin (Mon; 10mg/kg, i.p, 3x/week) treatment in immunocompromised mice bearing SK-MEL-28 shControl or shp16 tumors. **(B-C)** Body weight of mice bearing **(B)** YUMM5.2 (mouse) or **(C)** SK-MEL-28 (human) tumors over the course of vehicle or Monensin treatment. Mean ± SD; one-way ANOVA.

## ACKNOWLEDGEMENTS

We thank The Wistar Institute Flow Cytometry Facility, Animal Facility, Genomics Facility, Bioinformatics facility, and Histotechnology Facility for their invaluable assistance with this project. Funding support for The Wistar Institute core facilities was provided by Cancer Center Support Grant P30CA010815. We also utilized resources from the Hillman Cancer Center shared facilities, partially supported by award P30CA047904, including the UPMC Hillman Cancer Center Cancer Genomics Core (RRID:SCR_025357) and Cancer Bioinformatics Services (RRID:SCR_025355), which supported analysis of the CRISPR screening data. Cancer Bioinformatics Services were additionally supported by the University of Pittsburgh Center for Research Computing, including the HTC high-performance computing cluster supported by NIH (S10OD028483). We also extend our gratitude to current and former members of the Aird Lab—Dr. Miho Naruse, Dr. Naveen Kumar Tangudu, Jeff Danielson, and Hui Wang —for their critical feedback throughout the project and manuscript preparation. Finally, we thank Francisco J. Vazquez Moreno for his exceptional contribution to the graphic illustrations included in this work.

## AUTHOR CONTRIBUTIONS

Amandine Amalric: Conceptualization, Investigation, Methodology, Visualization, Writing – Review & Editing. Baixue Yang: Conceptualization, Investigation, Methodology, Visualization, Writing – Review & Editing. Rebecca Scarpati: Investigation, Methodology, Visualization. Apoorva Uboveja: Investigation, Writing – Review & Editing. Ruiying Zheng: Investigation, Methodology, Visualization. Alex Tom: Investigation, Methodology, Visualization. Aidan R. Cole: Investigation, Writing – Writing – Review & Editing. Andrea Andress Huacachino: Investigation, Methodology, Visualization. Qin Liu: Statistical Support, Writing – Review & Editing. Nathaniel W. Snyder: Investigation, Methodology, Visualization, Writing – Writing – Review & Editing. Andrew V. Kossenkov: Investigation, Writing – Review & Editing. Katherine M. Aird: Conceptualization, Investigation, Visualization, Writing – Review & Editing, Supervision, Project Administration, Funding Acquisition. Raquel Buj: Conceptualization, Investigation, Methodology, Visualization, Writing – Original Draft, Review & Editing, Supervision, Funding acquisition. All authors reviewed the results and approved the final version of the paper.

## MATERIAL AND METHODS

### Cell Lines and culture conditions

All cell lines used in this study were purchased from ATCC and were used within 50 passages. ATCC performs cell line authentication by short tandem repeat (STR) profiling. Cells were cultured within 3 months of receipt or resuscitation. The murine melanoma cell lines YUMM5.2 (ATCC, catalog no. CRL-3367, RRID: CVCL_JK43) were cultured in DMEM and Ham’s F-12 50/50 media (Corning, catalog no. 10-092-CV), supplemented with nonessential amino acids (Corning, catalog no. 25-025-Cl) and 10% FBS (BioWest, catalog no. S1620). The human melanoma cell line SK-MEL-28 (ATCC, catalog no. HTB-72, RRID: CVCL_0526), and the human embryonic kidney cell line HEK293-FT (ATCC, catalog no. CRL-3216, RRID: CVCL_6911) were cultured in DMEM (Corning, catalog no. 10-013-CV), supplemented with 5% FBS (BioWest, catalog no. S1620). All cell lines were supplemented with 1% Penicillin/Streptomycin (Gibco, catalog no. 15-140-122) and maintained at 37°C with 5% CO2. To ensure the absence of Mycoplasma contamination, all cell lines were tested monthly as described in^58^.

### Plasmid information

To perform knockdown experiments, short hairpin RNAs (shRNAs) targeting specific genes were used. The shRNA sequences employed in this study included: human shp16 (TRCN0000010482) and mouse sh*Cdkn2a* #1 (TRCN0000077816) and #2 (TRCN0000362595) were purchased pre-cloned into the lentiviral pLKO.1 vector (Sigma-Aldrich). Control lentiviral vector pLKO.1-Puro shGFP (Addgene, catalog no. 30323, RRID: Addgene_30323), was used for both human and mouse experiments.

### Lentiviral Packaging and Infection conditions

Lentiviral vectors were packaged as follows: 10 µg of the corresponding lentiviral vector was mixed with 7.5 µg of psPAX2 (Addgene, catalog no. 12260, RRID: Addgene_12260) and 7.5 µg of pMD2.G (Addgene, catalog no. 12259, RRID: Addgene_12259) in 1.5 mL of Opti-MEM™ media (Gibco, catalog no. 31985070) and incubated for 5 minutes at room temperature. In parallel, 50 µL of Lipofectamine™ 2000 Transfection Reagent (Invitrogen, catalog no. 11668019) was diluted in 1.5 mL of Opti-MEM™ media and incubated for 5 minutes at room temperature. The Lipofectamine mixture was added dropwise to the plasmid mixture, gently swirled to mix, and incubated for 20 minutes at room temperature. The Lipofectamine-lentiviral complexes were then added dropwise to a 60-70% confluent 10 cm dish of HEK293-FT cells, seeded 24 hours in advance and containing 7 mL of regular media, and incubated at 37°C with 5% CO_2_. Six to eight hours later, the media was replaced with 9 mL of fresh regular media and incubated at 37°C with 5% CO_2_. Seventy-two hours later, the viral supernatants were collected, filtered through a 45 µm filter (Fisherbrand™, catalog no. 09-720-005), and used to infect target cells for 16 hours. Infected cells were selected for 3 days with puromycin at 1 µg/mL for SK-MEL-28 cells or 3 µg/mL for YUMM5.2 cells (Thermo Fisher Scientific, catalog no. A1113802).

### CRISPR Drop out Screen

The mouse metabolic-focused CRISPR knockout library screen performed in YUMM5.2 mouse melanoma cells with stable knockdown of Cdkn2a (shCdkn2a) or shGFP control was previously published by our group, and additional details regarding library composition, screening conditions, sequencing, and primary analysis are provided in the original publication^7^. For the present study, we reanalyzed this existing dataset using a distinct analytical framework aimed at identifying higher-order cellular dependencies associated with p16 loss rather than individual gene-level hits. Specifically, genes that were significantly depleted in shCdkn2a cells relative to shGFP controls, defined as genes targeted by more than five sgRNAs with p < 0.05, were classified as essential for survival in the p16-deficient context (**Table S1**). This gene set was then used as input for Gene Set Enrichment Analysis (GSEA) focusing on Gene Ontology Cellular Component (GOCC) annotations to identify organelle-level vulnerabilities.

### *In vitro* treatments

Equal numbers of cells were seeded and allowed to adhere for 24h prior to treatment. Cells were then treated with Golgi-disrupting agents, including Monensin (Millipore Sigma, catalog no. M5273; 0.12µM for YUMM5.2 and SK-MEL-28 cells), Brefeldin A (Millipore Sigma, catalog no. B7651-5MG; 405nM for YUMM5.2 and 45nM for SK-MEL-28 cells), or Golgicide A (Millipore Sigma, catalog no. G0923-5MG; 6µM for YUMM5.2 and 0.6µM for SK-MEL-28 cells). Pharmacologic induction of lipid peroxidation was achieved using RSL3 (Millipore Sigma, catalog no. SML2234-5MG; 5µM for YUMM5.2 and SKMEL-28 cells, for 6h). Where indicated, cells were co-treated with the lipid peroxidation inhibitor Liproxstatin-1 (Selleck Chemicals, catalog no. S7699; 1µM for YUMM5.2 and SK-MEL-28 cells). All treatments were performed in triplicate and maintained for 4 days, after which cells were collected for downstream analyses.

### Western Blotting

Cell lysates were collected in 1X sample buffer [2% SDS (Millipore Sigma, catalog no. 75746-1Kg), 10% glycerol (Sigma Aldrich, catalog no. G5516), 0.01% bromophenol blue (Millipore Sigma, catalog no. BX1410), 62.5 mM Tris (Fisher Scientific, catalog no. AAJ65594A1), pH=6.8 and 0.1 M DTT (Thermo Fisher, catalog no. 15508-013)] and boiled at 95° C for 10 minutes. Protein concentration was determined using Bradford assay (Bio-Rad, catalog no. 5000006) following manufacturer’s instructions. Proteins were resolved using SDS-PAGE gels and transferred to nitrocellulose membranes (GE Healthcare Life Sciences, catalog no. 10600001). Antibodies used include: anti-human p16 (Abcam, catalog no. ab108349, working concentration 1.5 µg/mL), anti-mouse p16 (Cell Signaling Technology, catalog no. 29271S, working concentration 1 µg/mL), anti-β-Actin (Sigma-Aldrich, catalog no. A1978, working concentration 1 µg/mL), anti GM130 antibody (BD Biosciences; catalog no. 610823; working concentration 1 µg/mL), anti-mouse HRP (Cell Signaling Technology, catalog no. 7076, working concentration 0.1 µg/mL), and anti-rabbit HRP (Cell Signaling Technology, catalog no. 7074, working concentration 0.2 µg/mL).

### Immunofluorescence assay and quantification

Thirty thousand cells were seeded onto coverslips and incubated at 37°C with 5% CO₂. Approximately 16 hours after seeding, the cells were fixed with 4% paraformaldehyde (Sigma Aldrich, catalog no. 158127-3Kg) for 10 minutes. Cells were washed out three times with PBS and permeabilized with 0.2% Triton X-100 in PBS for 5 minutes. Cells were blocked for 5 minutes with 3% BSA/PBS followed by incubation with 1/200 dilution of anti GM130 antibody (BD Biosciences; catalog no. 610823; RRID:AB_398142) in 3% BSA/PBS for 1 hour at room temperature. Cells were washed three times with 1% Triton X-100 and incubated with Cy3 conjugated secondary antibody (Jackson Immunoresearch, catalog no. 712-165-150; RRID:) diluted 1/1,000 in 3% BSA/PBS and incubated 1 hour at room temperature and in dark. Cells were stained with 0.15 μg/mL DAPI (diluted in PBS) for 1 minute and washed three times with PBS. Finally, coverslips were mounted (Sigma Aldrich, catalog no. F6182-20ML) and sealed with transparent nail polish. Controls included staining with 10 µg/mL normal rabbit IgG (Cell Signaling, catalog no. 2729) and secondary-only staining. At least 200 cells per coverslip were counted. Cells were differentiated into 3 categories based on the morphology of Golgi apparatus: perinuclear, compacted (Golgi apparatus are localized in juxtanuclear area) and dispersed.

### Dose-response assays

Equal numbers of cells were seeded and allowed to adhere for 24h prior to treatment. Cells were then treated for 4 days with Golgi-disrupting agents, including Monensin (up to 30 µM), Brefeldin A (up to 25 µM), or Golgicide A (up to 10 µM), either alone or in combination with Liproxstatin-1 (1 µM). Following treatment, cell viability was assessed by fixing cells with 1% paraformaldehyde for 5 min and staining with 0.5% crystal violet. Excess stain was removed by washing with tap water. After drying for 2-3 days, bound dye was solubilized with 10% acetic acid, and absorbance was measured at 590 nm using a BioTek Epoch microplate reader. All conditions were performed in sextuplicate.

### BODIPY 581/591 C11 staining and flow cytometry analysis

A total of 120,000 SK-MEL-28 shControl and shp16 cells and 80,000 Yumm5.2 shControl and sh*Cdkn2a* (hairpins #1 and #2) cells were seeded in triplicate in 6-well plates and incubated for 4 days. Cells were then trypsinized and incubated with 5 µM BODIPY C11 (ThermoFisher Scientific, catalog no. D3861) in 10% FBS/DPBS for 10 minutes at 37°C. Cells were subsequently washed 1 time with DPBS and put into 2% FBS/DPBS solution. Lipid peroxidation was analyzed by flow cytometry (BD Symphony A3). Data analysis was performed using FlowJo software (version 10.10.1).

### Mouse models and treatments

All mice were housed in a HEPA-filtered ventilated rack system at the Animal Facility of The Wistar Institute and maintained on a 12-hour light/dark cycle at 21 ± 2°C. Mice were group-housed with up to five animals per cage. All animal procedures were approved by the Institutional Animal Care and Use Committee (IACUC) of The Wistar Institute. A total of 350,000 YUMM5.2 cells (shControl or shCdkn2a) were subcutaneously implanted into the right flank of 6–8-week-old male C57BL/6J mice (The Jackson Laboratory, strain #000664, RRID: IMSR_JAX:000664), and 10,000,000 SK-MEL-28 cells (shControl or shp16) were subcutaneously implanted into the right flank of 6–8-week-old male NOD.Cg-Prkdc^scid^ Il2rg^tm1Wjl^/SzJ mice (The Jackson Laboratory, strain #005557, RRID: IMSR_JAX:005557). Mice were monitored daily until tumors became palpable, after which animals were randomly assigned to vehicle or monensin treatment groups. Mice were treated five days per week by intraperitoneal injection with monensin (10 mg/kg; Millipore Sigma, catalog no. M5270) diluted in 200 µL PBS, or with equal volume of vehicle. Body weight and tumor size were measured three times per week. Tumor volume was calculated using the formula ½ × (Length × Wide²) where Length>Wide. The experimental endpoint for longitudinal tumor growth studies was defined as the time point when the first mouse reached the IACUC-approved tumor volume criterion of >1 cm³. For immunohistochemistry (IHC) studies, the endpoint was defined when the mean tumor volume of the cohort reached 0.1 cm³.

### Immunohistochemistry (IHC)

IHC was done as described previously^59^. After deparaffinization and rehydration, tissues were steamed for 40 min in 1x citrate buffer (Sigma-Aldrich, catalog no. C9999), after which they were immersed in 3% MeOH for 20 min. Tissues were blocked with 1% BSA in PBS for 30 min before overnight incubation using normal rabbit IgG (2µg/ml, Cell Signaling Technology, catalog no. 2729S) and anti-4-HNE (2µg/ml, R&D Systems, catalog no. MAB-3249). A mouse- and rabbit-specific HRP/DAB (ABC) detection IHC kit (Abcam, catalog no. 4264) was used for visualization of staining. Tissues were incubated in Mayer’s haematoxylin solution (Sigma-Aldrich, catalog no. MHS16) for 3 min followed by bluing in running tap water for 2 min. Tissues were dehydrated and mounted in Cytoseal (Sigma-Aldrich, catalog no. C23-244257). Data analysis was performed using QuPath software (version 0.6.0). Due to high background, 4-HNE H-score was calculated by subtracting each tissue’s IgG H-score.

### RNA-Seq

Total RNA was extracted from cells with Trizol (Ambion, catalog no. 15596018) and DNase treated, cleaned, and concentrated using Zymo columns (Zymo Research, catalog no. R1013) following the manufacturer’s instructions. RNA integrity number (RIN) was measured using RNA Screen Tape and TapeStation Analysis Softawre (version 5.1) to confirm a RIN greater than 7 for each sample. The cDNA libraries, next-generation sequencing, and bioinformatics analysis were performed by The Wistar Genomics facility, Azenta and The Wistar Bioinformatics Facility, respectively. Differential expression analysis was done using R CRAN (version 4.6.1) and DESeq (version 1.52.0). Genes were considered differentially expressed if they had an adjusted p value < 0.05 and a log₂ fold change (log₂FC) ≥ 1.5 for upregulated genes or ≤ −1.5 for downregulated genes.

For volcano plots (**Fig. S1D-E**), genes associated with the gene sets Golgi Cisterna Membrane and Golgi Cis Cisterna (GO CC) were obtained from the GSEA resource. Likewise, for heatmaps (**Fig. S4C-D**), genes associated with the Ferroptosis pathway (WikiPathways) and the Antioxidant Activity gene (GO MF) were obtained from the GSEA resource. Species-specific gene sets were used for the murine YUMM5.2 and human SK-MEL-28 cell lines. Only genes successfully detected in the respective RNA-seq datasets were included in the plots. For Venn diagram analyses (**Fig. S3**), differentially expressed genes were identified using the criteria described above. In the YUMM5.2 and SK-MEL-28 models, only genes significantly differentially expressed following monensin treatment relative to DMSO in sh*Cdkn2a* or shp16 cells, respectively, that were not significantly differentially expressed in the corresponding shControl cells were included. For THP-1 cells, raw RNA-seq data from GSE130657^34^ were obtained from the GEO, and differential expression analysis comparing monensin-treated and DMSO-treated cells was performed using DESeq2. Differentially expressed genes were defined using the same adjusted p-value and log_2_FC thresholds. For A549 cells, differential expression results were obtained from the published study^35^, which reported differentially expressed genes based on an adjusted p-value < 0.05 following 8-hour monensin treatment. Genes meeting the log_2_FC threshold of ≥ 1.5 or ≤ −1.5 were included in the analysis. Human and mouse gene identifiers were mapped to their corresponding orthologous gene identifiers using Ensembl BioMart.

### Qualification and Statistical analysis

GraphPad Prism version 10.0 was used to perform statistical analysis. The level of significance between two groups was assessed with unpaired t test. For data sets with more than two groups, one-way ANOVA followed by Šídák’s post hoc test was applied. P-values< 0.05 were considered significant.

